# PanGu Drug Model: Learn a Molecule Like a Human

**DOI:** 10.1101/2022.03.31.485886

**Authors:** Xinyuan Lin, Chi Xu, Zhaoping Xiong, Xinfeng Zhang, Ningxi Ni, Bolin Ni, Jianlong Chang, Ruiqing Pan, Zidong Wang, Fan Yu, Qi Tian, Hualiang Jiang, Mingyue Zheng, Nan Qiao

**Author notes:** Authors contribute equally (Xinyuan Lin, Chi Xu).

## Abstract

Recent achievements in large-scale pre-trained models like GPT-3 and PanGu-α showed amazing performances in many downstream tasks, which makes AI friendlier toward industrial users. Deep learning has been recognized as the most promising technology for pharmaceuticals, a powerful molecule pre-trained model could save researchers tons of time. In chemistry classes, the students learn two molecule representations, the molecular formula and the structure formula, and learn to translate them from one way to the other. Inspired by this, we developed a novel deep learning architecture using a graph-to-sequence asymmetric conditional variational autoencoders, called PanGu Drug Model, which can appropriately characterize the molecule from both representations and improve the performance of downstream drug discovery tasks. After pretrained with 1.7 billion small molecules, our proposed model achieved the state-of-the-art results in 20 drug discovery tasks, such as molecule property prediction (predict ADMET properties, compound-target interactions, drug-drug interactions and chemical reaction productivity), molecule generation and molecule optimization. A new drug screening library of 100 million drug-like small molecules with 99.68% novelty was generated by PanGu molecule generator, which could efficiently produce novel compounds with similar physiochemical properties to given distribution, this library could be used to supplement existing compound databases. In addition, PanGu molecule optimizer could optimize the chemical structures of starting molecule with improved molecular property of interest. An automatic multi-objective optimization web application implemented by PanGu Drug Model is available at http://www.pangu-drug.com/.

## Introduction

Facing the situation that more than thousands of diseases exist whereas most of them have no cure treatments, for example many kinds of rare diseases, novel drug discovery plays a significant important role^1,2^. Besides, complicated and sudden illness endanger human beings’ life and health. Therefore, research and development (R&D) of new drugs is a necessary demand to the public. Traditional new drug R&D usually costs tens to hundreds of millions of U.S. dollars and more than ten years of time duration per drug from research to marketing^3^. In recent years, the rapid development of new experimental technologies in the field of drug R&D such as cryo-electron microscopy, DNA-encoded compound libraries, various omics technologies, etc., has produced massive amounts of data, which allows artificial intelligence (AI) technology to dig deeper into these data and offers a new paradigm for new drug R&D.

The concept of AI drug discovery (AIDD) began to be proposed in 2016, and has been verified in 2019, including: the AI cooperation between *Exscientia* and Japan’s *Sumitomo* Pharmaceutical entered the clinic and shorten the optimization process into only one year; Recursion Pharmaceuticals collaborated with Takeda Pharmaceutical found new drug candidates for six rare diseases, which took 1.5 years; in 2020, *Insilico Medicine* quickly discovered *DDR1* drug candidates, taking only 46 days from the establishment of the project to the discovery of the drug candidates, and this process usually takes several years^4,5^. These cases showed that AI can significantly improve the efficiency of drug discovery.

The core of drug discovery is through repeating the cycle of drug design, screening and optimization, to find effective compounds to targets, and meanwhile guarantee the safety standards to human beings. Besides those recently reported successful AIDD researches, there have been series of computer-aided drug discovery (CADD) methods, such as structure-based and ligand-based drug virtual screening for hit identification and quantitative structure-activity relationship (QSAR) for molecule property prediction^6^. However most of them only contributed in improvement of one step of drug R&D, a unified model capable of empowering the whole-process drug discovery still lacks. Possible reasons to the phenomenon are the few percentages of small molecules with golden standard labels: 1) Drug discovery tasks are various and complicated. Corresponding experimental data rarely share common molecules between tasks, which make it difficult to build a model to predict all the tasks simultaneously. 2) Some property records only cover a few numbers of chemicals so it lacks enough labelled data to train a model.

In contrast to the few labelled number, public databases already contain tens of billions of chemical structures of available small molecules without detailed annotations e.g. ZINC20^7^, ChEMBL^8^ and PubChem^9^ etc.. Efficiently integrating these large-scale chemicals with the labeled ones can benefit drug discovery study. Traditionally, Extended-Connectivity Fingerprints (ECFP)^10^ are circular topological fingerprints designed for molecular characterization, similarity searching, and structure-activity modeling, which have been popular among a wide variety of drug R&D applications. However, it consists of only 0 or 1 indicating presence or absence of certain functional structures, cannot continuously describe molecular characteristics. Additionally, due to the existence of bit collisions, ECFP sometimes represents different structures using the same bit features. And the fixed calculation rule of ECFP disables it adaptively updated on specific training data.

To make the representation continuously learnable from expanding chemicals, suitable to train on specific downstream tasks and construct a representation decoder applicable to generate novel molecule structures, we built a novel pre-training Conditional Variational Autoencoder (cVAE)^11^ based model, capable of exactly representing, compressing and reconstructing molecules’ chemical structures. Unlike classic autoencoders, we invented a novel graph-to-sequence (graph2seq) asymmetric architecture to achieve the above purposes, called PanGu Drug Model (PanGu for short, pronounce as pang-goo), and used 1.7 billion chemical structures to pre-train it, with input of chemicals’ 2D undirected cyclic graphs^12^ and output of corresponding formula strings. Here the procedures are like this - humans read an image of a chemical structure and write down a text of the corresponding chemical formula, thus after repeating this for billions of times PanGu can learn the human-like cognition transformation between chemical structures and formula strings. So we called this modeling idea as *Molecular Structure Formulating Transformation* (MSFT). Compared to MSFT, other types of inputs and outputs have the following disadvantages: 1) Sequence input cannot learn molecular information of the connections between atoms. 2) Graph output models require calculating reconstruction loss for generated graphs. The problem of graph isomorphism can be addressed by graph matching however requires expensive computation^13^.

In 2020, two large-scale pre-training models were published. GROVER^14^ is a graph representation model from message passing transformer, with self-supervised tasks in node-, edge- and graph-level; it contains about 100 million parameters and was pre-trained with 11 million small molecules. X-Mol^15^ is a transformer-based language model that train through input random Simplified Molecular Input Line Entry Specification (SMILES)^16^ and generate another valid one; it was pre-trained with 1.1 billion small molecules. Both GROVER and X-Mol were compared in our study.

Here, we use pre-trained PanGu for total 20 tasks covering all steps of drug screening process, and showing the state-of-the-art performance in 16 kinds of molecule property prediction (predicting compound-target interactions (CPIs), absorption, distribution, metabolism, excretion and toxicity (ADMET) scores, drug-drug interactions (DDIs) and chemical reaction productivity), molecule generation, constrained optimization, etc.

There are two ways to applying pre-trained PanGu on specific supervised learning tasks: 1) Fine-tuning of PanGu, by directly integrating the encoder of PanGu with a predictor, to make the encoder better capture the intrinsic pattern of the task; 2) Using *PanGu Fingerprint*, which is the combination of the latent space and the condition vector of cVAE, can be used as a type of representation for molecules (fixed-size and ECFP-like). While using PanGu Fingerprint, a simple multilayer perceptron (MLP) can be used as predictor which accepts a vector as molecule input and the parameters of pre-trained PanGu backbone remains unchanged during training.

Additionally, PanGu molecule generator efficiently produced novel chemicals with similar physiochemical properties to given distribution. Using it, a novel drug library of 100 million drug-like chemicals with 99.68% novelty were automatically generated. And, PanGu molecule optimizer could optimize the chemical structures of starting molecule with improved molecular property of interest. Furthermore, an automatic multi-objective optimization is implemented and can be freely accessed at http://www.pangu-drug.com/.

## Results and discussion

Computer-aided technology can accelerate candidate drug screening extremely quickly. We designed and trained PanGu by taking advantage of billions of chemicals to learn molecular representation and reconstruction. Besides, seven experiments with totally twenty tasks were conducted by using the pre-trained model, which cover almost all aspects of drug screening scenarios (Fig. 1).

**Fig. 1.**
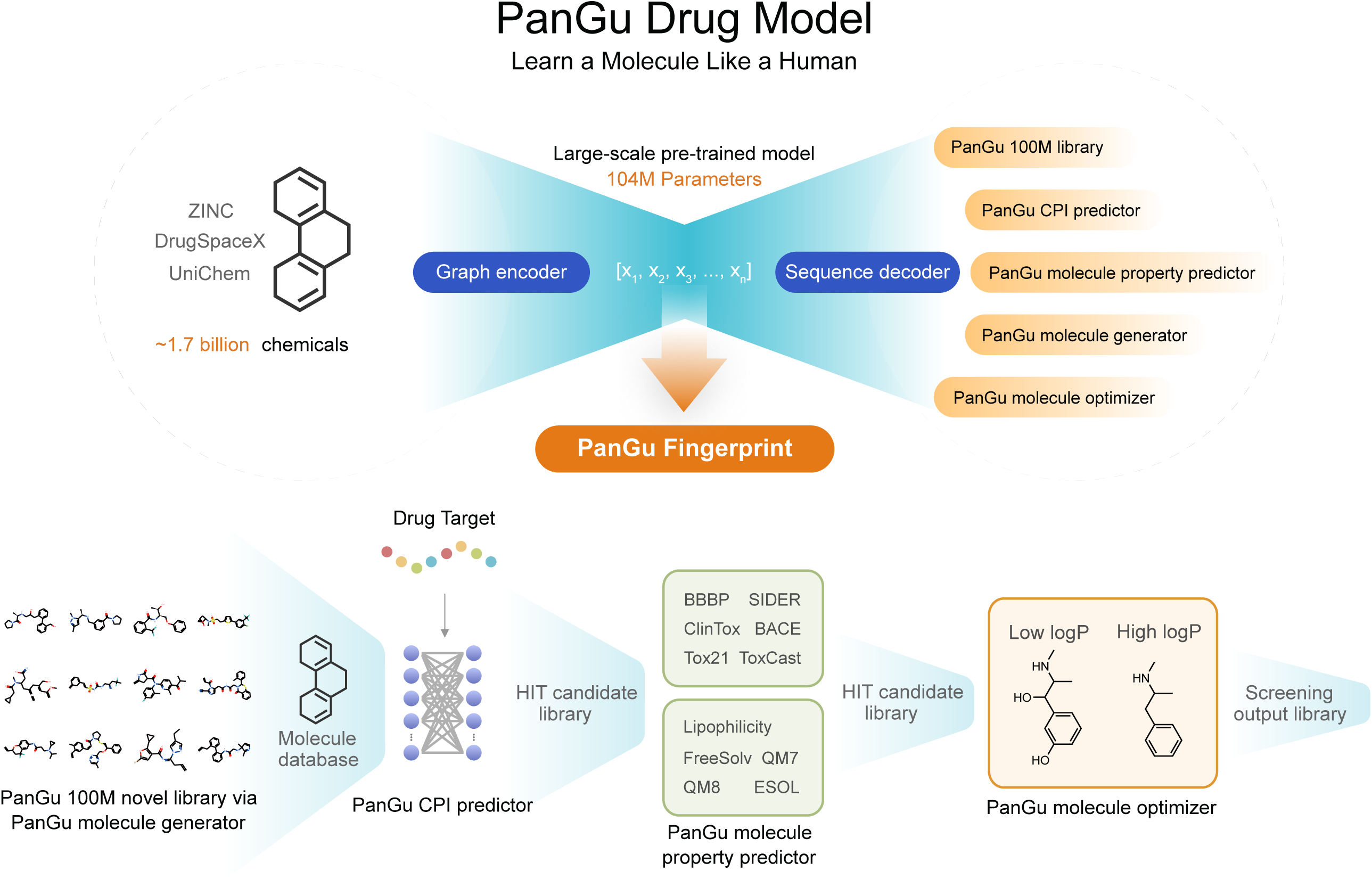
PanGu for AI drug screening. Schematic diagram of the process of PanGu training and PanGu application to a variety of AI drug screening tasks. The upper shows the conditional variational autoencoder structure of PanGu, pre-trained with chemical structures of 1.7 billion small molecules. The below shows the AI-powered drug screening flowchart consisting of AI tasks of compound-protein interaction prediction, molecular property prediction, drug-like optimization and library generation, and shows the molecule screening procedure from an initial molecular database, to a HIT candidate library, lead-like library and final screening output library. Arrows indicate PanGu Fingerprint and PanGu decoder empower the corresponding tasks.

### Molecule property prediction

Molecular property evaluation plays an important role in lead compound selection, especially ADMET properties and bioactivity to certain targets^17^. Here, we totally tested PanGu on 11 molecule property prediction tasks (see the Detailed information about molecule property prediction tasks part of the Supplementary Information section) from MoleculeNet^18^. Here, fine-tuned PanGu and PanGu Fingerprint were both evaluated. For fine-tuning, the encoder was followed by a multilayer perceptron predictor, and parameters of encoder were updated along with the predictor under a low learning rate for each task. The fine-tuned PanGu was compared to GROVER^14^, GCN^19^, Weave^12^, SchNet^20^, MPNN^21^, DMPNN^22^ and AttentiveFP^23^. And for PanGu Fingerprint, ECFP6 with 2048-bit length (same length as ours PanGu Fingerprint) which is obtained by RDKit^24^ was compared as baseline. We followed the same training and evaluating strategies as GROVER (Rong et al., 2020) and reported the performance on test set. It is worth mentioning that, GROVER employed 200-dimensional 2D features obtained by RDKit to achieve competitive performance while PanGu didn’t.

The results showed that fine-tuned PanGu outperformed existing deep learning models in all the property prediction tasks, with a better average or lower standard derivation in performance (Table 1). Compared to most recently published pre-training model GROVER, PanGu obtained about 0.8% average improvement in accuracies of classification tasks, and errors (root mean square error (RMSE) or mean absolute error (MAE)) of regression tasks were decreased about 4.9%. For example, in the task of ESOL, which is predicting the solubility of compounds, PanGu got the average RMSE of 0.747 which is 10.1% lower than 0.831 of GROVER. In order to compare the performance improvement under different data scales, we divided the 11 tasks into larger-data and smaller-data groups based on number of molecules within (2000 as threshold). Interestingly, we found that PanGu showed better improvement for the smaller-data tasks (+1.2% and +0.4% for accuracy improvement of classification tasks with smaller dataset and larger dataset respectively, and -8.5% and -2.4% for error decrement of regression tasks, respectively).

**Table 1.**
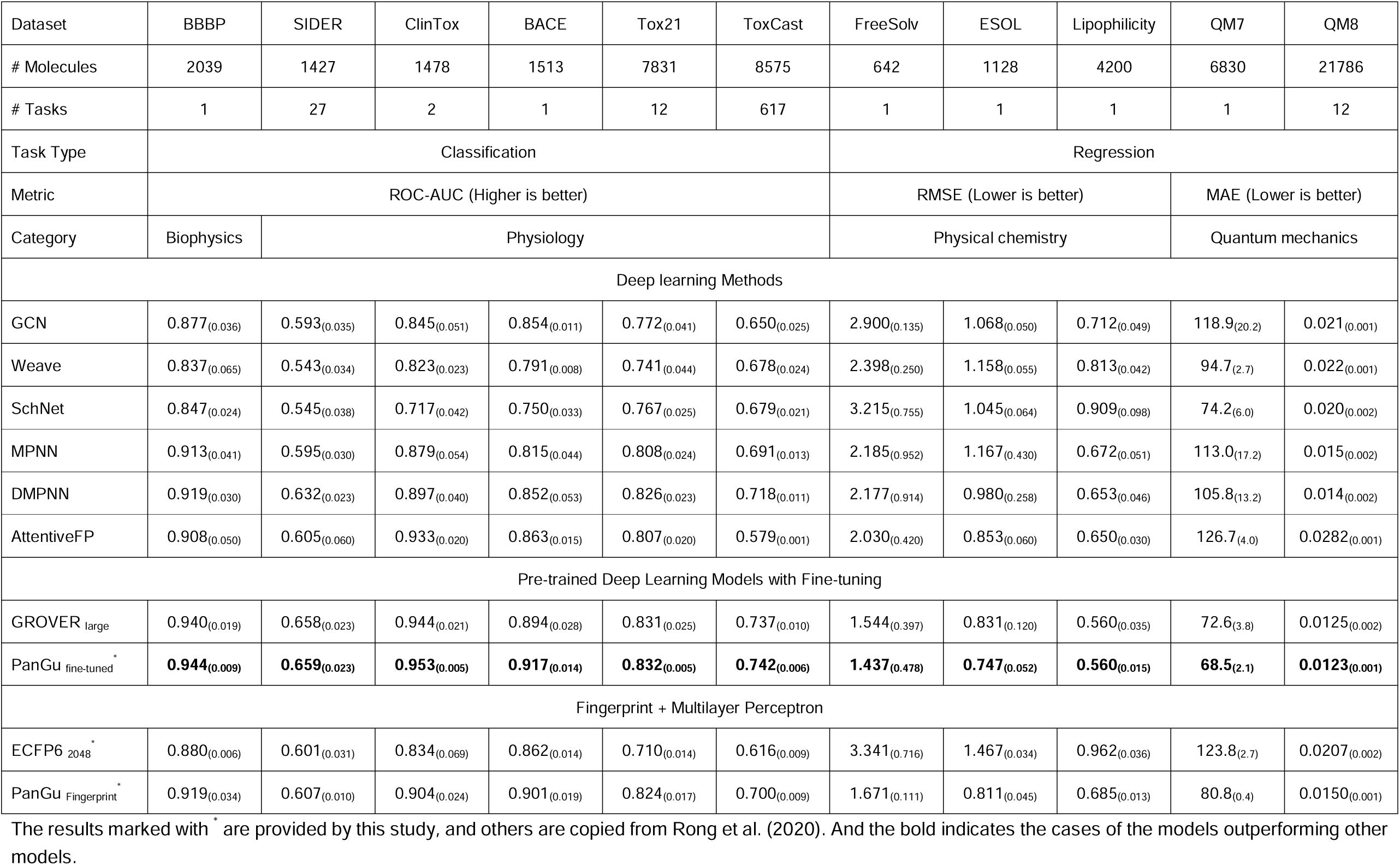
Molecule property prediction results. Mean and standard deviation of 3 independent trials with randomly seeded scaffold splitting.

For PanGu Fingerprint, it outperformed ECFP on all tasks, with about +8.0% of average improvement in classification accuracy and about 37.2% of average decrease in regression error. Surprisingly, PanGu Fingerprint obtained better results than several deep learning models especially in following 7 tasks: BACE, BBBP, Tox21, ToxCast, FreeSolv, ESOL and QM7. In tasks of BACE and ESOL, PanGu Fingerprint even outperformed GROVER and ranked the second place, only after the fine-tuned PanGu.

The result of molecule prediction tasks confirmed the expressive ability of PanGu in both fine-tuned model and fingerprint, which indicates that PanGu can provide more effective guidance in the drug screening process. It is noteworthy that, except the huge improvement to ECFP, using PanGu Fingerprint for prediction can greatly reduce the computational overhead compared to fine-tuning strategy. Therefore, it can be considered as an alternative to ECFP in most cases of molecule-related prediction. Besides, we believe that another advantage of PanGu Fingerprint compared to other VAE-based models is the design of layered latent space (see the Graph-to-sequence architecture of PanGu part of Methods section, and the Ablation studies of layered latent space of the Supplementary Information section), which describing molecules at different radius scales and chemically interpretable.

### Compound-protein interaction prediction

Compound-protein interaction (CPI) identification lies in the first procedure of drug screening to search for active hit compounds against given protein targets^25^. Here, we evaluated PanGu on three previous benchmark datasets, human dataset^26,27^, *C. elegans* dataset^27^ and BindingDB dataset^28^. Area Under Receiver Operating Characteristic Curve (ROC-AUC), Area Under Precision Recall Curve (PRC-AUC), precision and recall were reported as previous studies did (see the Detailed information of experiments part of the Methods section). We included four relevant published deep learning models TransformerCPI^29^, GraphDTA^30^, GCN^19^, CPI-GNN^27^ and four machine learning methods i.e. K nearest neighbors (KNN), L2-logistic (L2) and random forest for comparison. The training and evaluating strategies were performed in accordance with Chen et al.’s study to ensure a consistent comparison (see the Detailed information of experiments part of the Methods section).

Consistently, in both human, *C. elegans* and BindingDB datasets (Table 2, Table 3 and Table 4), PanGu Fingerprint ranked first in the ROC-AUC, precision, recall and PRC, outperforming the seconding-ranking TransformerCPI (human data: ROC-AUC 0.985 vs. 0.973, precision 0.951 vs. 0.916, recall 0936 vs. 0.925; *C. elegans* data: ROC-AUC 0.994 vs. 0.988, precision 0.976 vs. 0.952, recall 0962 vs. 0.953; BindingDB data: ROC-AUC 0.968 vs. 0.951, PRC-AUC 0.969 vs. 0.949). This result showed that in addition to good performance in predicting molecule property, PanGu Fingerprint can also provide effective information when combined with other representations.

**Table 2.**
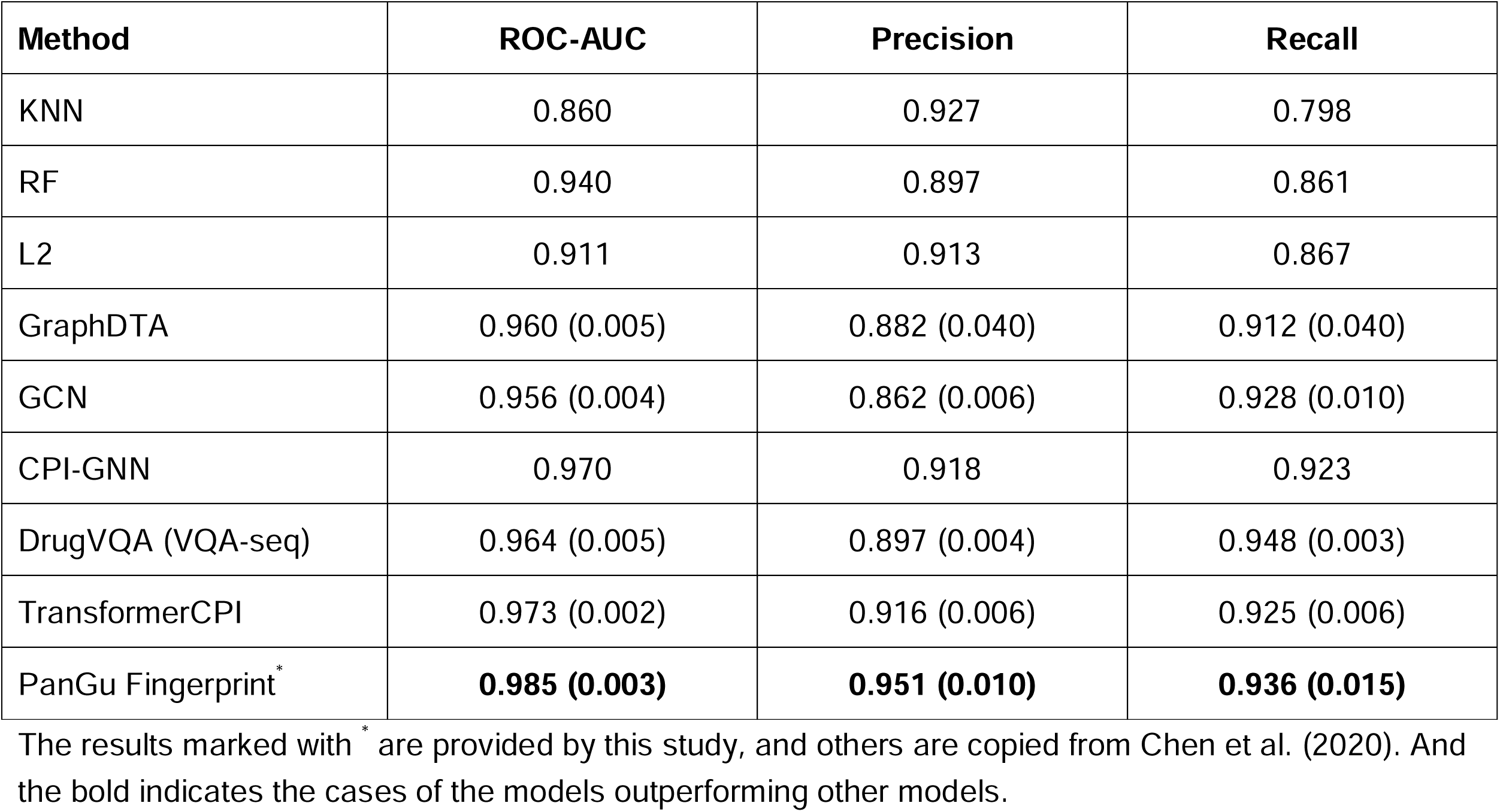
Compound-protein interaction prediction results on human dataset.

| Method | ROC-AUC | Precision | Recall |
| --- | --- | --- | --- |
| KNN | 0.860 | 0.927 | 0.798 |
| RF | 0.940 | 0.897 | 0.861 |
| L2 | 0.911 | 0.913 | 0.867 |
| GraphDTA | 0.960 (0.005) | 0.882 (0.040) | 0.912 (0.040) |
| GCN | 0.956 (0.004) | 0.862 (0.006) | 0.928 (0.010) |
| CPI-GNN | 0.970 | 0.918 | 0.923 |
| DrugVQA (VQA-seq) | 0.964 (0.005) | 0.897 (0.004) | 0.948 (0.003) |
| TransformerCPI | 0.973 (0.002) | 0.916 (0.006) | 0.925 (0.006) |
| PanGu Fingerprint<sup>*</sup> | **0.985 (0.003)** | **0.951 (0.010)** | **0.936 (0.015)** |

The results marked with <sup>\*</sup> are provided by this study, and others are copied from Chen et al. (2020). And the bold indicates the cases of the models outperforming other models.

**Table 3.** Compound-protein interaction prediction results on *C. elegans* dataset.

| Method | ROC-AUC | Precision | Recall |
| --- | --- | --- | --- |
| KNN | 0.858 | 0.801 | 0.827 |
| RF | 0.902 | 0.821 | 0.844 |
| L2 | 0.892 | 0.890 | 0.877 |
| GraphDTA | 0.974 (0.004) | 0.927 (0.015) | 0.912 (0.023) |
| GCN | 0.975 (0.004) | 0.921 (0.008) | 0.927 (0.006) |
| CPI-GNN | 0.978 | 0.938 | 0.929 |
| TransformerCPI | 0.988 (0.002) | 0.952 (0.006) | 0.953 (0.005) |
| PanGu Fingerprint<sup>*</sup> | **0.994 (0.001)** | **0.976 (0.009)** | **0.962 (0.005)** |

The results marked with <sup>\*</sup> are provided by this study, and others are copied from Chen et al. (2020). And the bold indicates the cases of the models outperforming other models.

**Table 4.** Compound-protein interaction prediction results on BindingDB dataset.

| Method | ROC-AUC | PRC-AUC |
| --- | --- | --- |
| GraphDTA | 0.929 | 0.917 |
| GCN | 0.927 | 0.913 |
| CPI-GNN | 0.603 | 0.543 |
| TransformerCPI | 0.951 | 0.949 |
| PanGu Fingerprint<sup>*</sup> | **0.968** | **0.969** |

The results marked with <sup>\*</sup> are provided by this study, and others are copied from Chen et al. (2020). And the bold indicates the cases of the models outperforming other models.

### Large-scale generative database

Molecule databases are indispensable for drug screening. Although many existing public databases are commonly used e.g. ZINC20^7^, ChEMBL^8^ etc., they only cover a small fraction of the entire chemical space, which restricts the screened drug to a known and limited chemical space^31^. Deep learning based newly generative chemicals can be used as data sources to search for effective compounds having not been recorded before. Generative models can learn the internal distribution of data during training, and then perform random sampling from the learned distribution to obtain new samples. And VAE-based models are capable of generation through decoding latent spaces. In order to supplement the existing drug databases with new chemical structures, a large-scale molecular generation task was designed, which aimed to generate 100 million molecules and evaluate the generated molecules with 3 statistical measurements, i.e. validity, uniqueness and novelty (see the Detailed information of experiments of the Methods section for more details). Here, any generated molecule appearing in ZINC20^7^, DrugSpaceX^31^ and UniChem^32^ databases were not considered as novel. Since there has been no deep learning study that produced such a large *in silico* dataset yet, we cannot directly compare the result to any existing baselines. Therefore, a small-scale molecule generation which adopting the same strategy of existing studies was performed and compared fairly (see the Additional experiments part of the Supplementary Information section for more details).

It showed that PanGu achieved a validity of nearly 100% (only 1,198 molecules of the 100 million were invalid), a uniqueness of 99.95% and a novelty of 99.68% (Table 5). Considering the large size of these databases, 100,000 samples were randomly selected from each set for analysis of chemical properties. We compared the distribution of chemical properties between the generated molecules and the prior set (Fig. 2). Interestingly, the generated molecules showed similar patterns to the prior set (most similar to ZINC due to it occupied 85% of the dataset). The t-Distributed Stochastic Neighbor Embedding (t-SNE) results (Fig. 2h) also showed that PanGu can generate molecules within a given chemical space quite well, but still with a high novelty. It demonstrates that PanGu have a good exploration ability to search inside the specific chemical space rather than simply to remember the training data. Besides, random 150 examples of generated molecules were showed in Supplementary Figure 5.

**Table 5.**
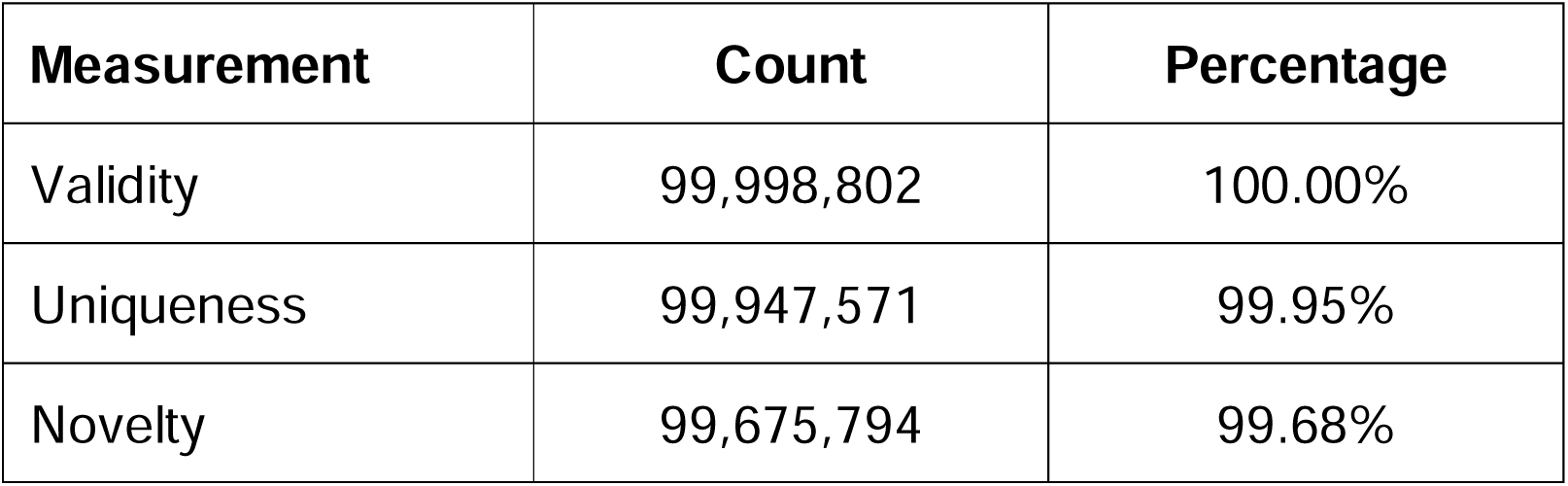
Large-scale molecule generation results.

| Measurement | Count | Percentage |
| --- | --- | --- |
| Validity | 99,998,802 | 100.00% |
| Uniqueness | 99,947,571 | 99.95% |
| Novelty | 99,675,794 | 99.68% |

**Fig. 2.**
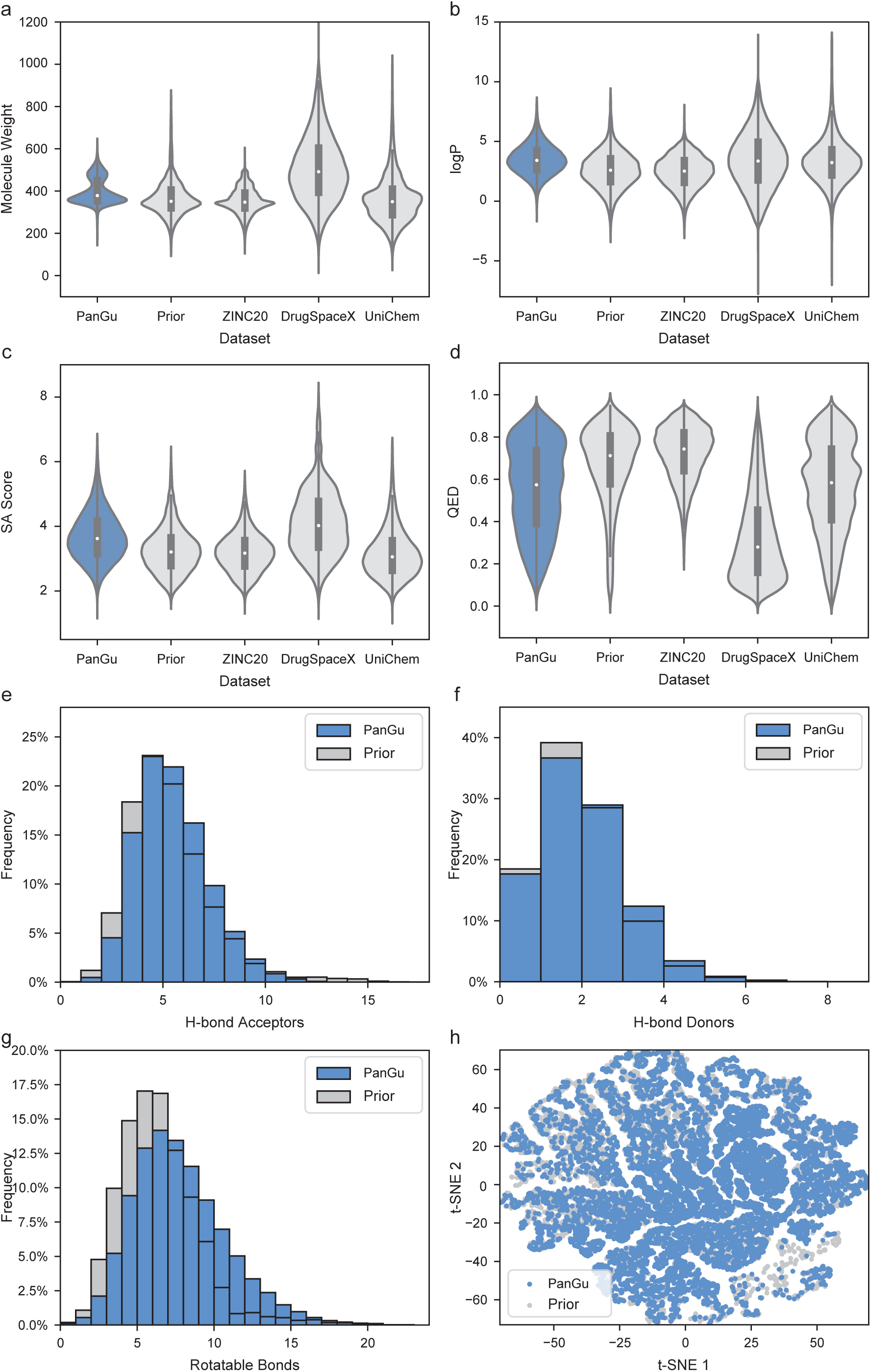
Comparative analysis of chemical properties of PanGu generated molecules to prior and existing public datasets. **a-d**, Violin plots of chemical properties of **a**, molecular weight, **b**, logP, **c**, SA, and **d**, QED of molecules generated by PanGu, the prior set for PanGu generation, ZINC20, DrugSpaceX and UniChem. logP is short for lipophilicity, where a higher value indicates more lipophilic. SA is short for synthetic accessibility, where a higher value indicates harder to synthesis. QED is short for quantitative estimation of drug-likeness, where a higher value indicates more drug-like. **e-g**, Distribution of numbers of **e**, H-bond acceptors, **f**, H-bond donors and **g**, rotatable bonds of molecules generated by PanGu, the prior set. **h**, A t-SNE plot comparing the chemical space of PanGu generated molecules and the prior set.

Generally, sequence-based generative models sometimes encounter the problem of low validity due to the syntax error^33^, and graph-based generative models are considered to have higher complexity and hard to train. PanGu leverages Self-Referencing Embedded Strings (SELFIES)^34^ to provide a simple and reliable way (see the Graph-to-sequence architecture of PanGu part of Methods section). The result of large-scale generation showed that, as long as we specify a distribution of desired molecules, the model can well explore and generate similar but novel molecules. That makes it possible to construct a large-scale deep learning generative drug-like molecular database. Furthermore, these generated novel molecules can be used for drug screening to enlarge the explored chemical space and enhance the drug screening task performance.

### Constrained molecule optimization

Development of new drugs usually starts from known and existing molecules^35^. Hit candidates from drug screening procedures are probably active for targets of interest but may not meet the needs in certain molecule properties. Thus, using PanGu we modified the hit candidate molecules to further improve the specific properties and meanwhile remain chemically similar to the original. Many deep learning models treat this task as a machine translation task, which translates the strings of SMILES from one to another with optimized properties. For PanGu, it is reasonable to assume that the latent space *z* containing the information of molecular structures and condition vector *c* controlling the molecular properties (see the Graph-to-sequence architecture of PanGu part of Methods section). By modifying conditional variables contained in condition vector *c*, we can directly achieve quantifiable and smooth optimization of the molecules for either single or multiple objectives. Here, molecule optimization was performed in a constrained scenario, and the penalized logP (the octanol-water partition coefficient (logP) penalized by the synthetic accessibility (SA) score and number of long cycles) was used as target chemical property. Given a molecule, the task was to find a different molecule *m’* that had the highest property value and remained a chemical structure similarity not less than a threshold *δ* to the, i.e. *sim*(*m, m*’) ≥ *δ* Here, we reported the optimized molecule with the highest property which satisfying the similarity constraint. If no molecules satisfy the constraint and has lower property than original one, the modification will be considered unsuccessful.

The results were shown in Table 6. When using four different thresholds *δ* of similarity to the initial molecules, PanGu all achieved higher improvement of the target property than JT-VAE. For example, when *δ* = 0.0 that is no similarity constraint, JT-VAE increased the target property by 1.91 on average, while PanGu significantly improved it by 9.13, about 4.8 times that of JT-VAE. Imaginably, the average improvement decreased along the similarity constraint *δ* increasing. Although when *δ* = 0.6, the average improvement of JT-VAE was only 0.21, PanGu still remained the optimization effect by 3.24, about 15.4 times that of JT-VAE.

**Table 6.** Constrained molecule optimization results on penalized logP. Mean and standard deviation of property improvement, molecular similarity and success rate under constraints *sim(m, m’) >= δ* with varied *δ*.

| | $\delta=0.0$ | | | $\delta=0.2$ | | |
| --- | --- | --- | --- | --- | --- | --- |
| Method | Improvement | Similarity | Success | Improvement | Similarity | Success |
| JT-VAE | 1.91 (2.04) | 0.28 (0.15) | 97.5% | 1.68 (1.85) | 0.33 (0.13) | 97.1% |
| PanGu | 9.13 (2.32) | 0.24 (0.12) | 100.0% | 8.34 (2.65) | 0.29 (0.11) | 100.0% |
| | $\delta=0.4$ | | | $\delta=0.6$ | | |
| Method | Improvement | Similarity | Success | Improvement | Similarity | Success |
| JT-VAE | 0.84 (1.45) | 0.51 (0.10) | 83.6% | 0.21 (0.71) | 0.69 (0.06) | 46.4% |
| PanGu | 4.89 (2.86) | 0.48 (0.09) | 91.8% | 3.24 (2.98) | 0.67 (0.08) | 48.9% |
| | $\delta=0.7$ | | | $\delta=0.8$ | | |
| JT-VAE | - | - | - | - | - | - |
| PanGu | 2.35 (2.75) | 0.79 (0.08) | 34.0% | 2.29 (2.89) | 0.87 (0.05) | 20.9% |

Baseline results are copied from Jin et al. (2018).

We chose four representative cases, which including two cases with high similarity to the starting molecule and other two cases with high target property improvement (Fig. 3). The first case had a similarity of 0.8462 and the penalized logP was improved by 2.542. Comparing the before- and after-optimized, we can find that two hydroxyl groups (OH) were removed by PanGu, which was in line with our chemical understanding (Fig. 3a). For the second case, interestingly PanGu replaced a certain side chain into a five-membered ring with an azanide, which increased the target property by 1.574 (Fig. 3b). The third one was a case with a property increase of 4.270 and a similarity of 0.4019. PanGu mainly optimized its adjacent ring (Fig. 3c), and removed the hydroxyl group. The forth one was a case with a large increase of 6.829 and with a similarity of 0.3725. It seems not similar between the two molecules by the first look, but PanGu turned the seven-membered ring with a hydroxyl group which adjacent to the five-membered ring into a benzene ring (highlighted in orange). And the other seven-membered ring was changed to a six-membered ring, and the hydroxyl group on it was also removed (highlighted in light blue). The side chain and the imidazole remained unchanged (Fig. 3d).

**Fig. 3.**
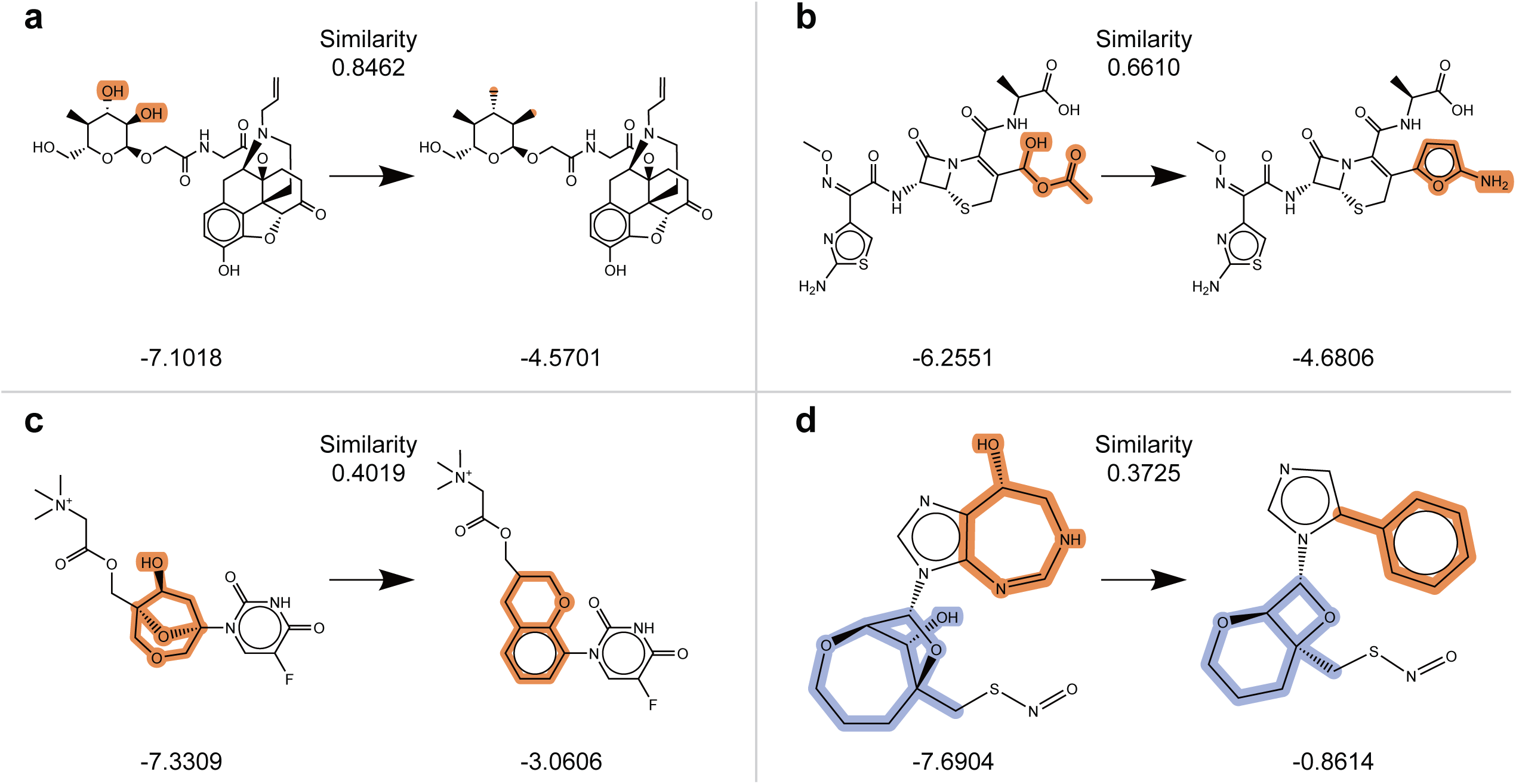
Cases of molecules optimized by PanGu for penalized logP. In each panel, left indicates the starting molecules, right indicates the optimized molecules by PanGu, corresponding penalized logP scores are shown below each molecule. Changes of molecules before- and after-optimization are highlighted, and the Tanimoto similarity scores between them are shown above.

We found that PanGu appears to be an efficient tool for molecule optimization by simply controlling the condition vectors. Compared to the baseline, PanGu greatly improved the target property of molecules while guaranteeing the optimization success rate but had only a slightly decrease in average similarity. This decrease seems acceptable while higher similarity to some extent is in conflict with greater property improvement. It is worth mentioning that, the number of long cycles (one of penalization terms) is not controlled conditionally in ours experiment but it did not affect the performance of our model. Besides, if we require other properties to be optimized, it is convenient to be integrated with PanGu for fine-tuning. Finally, a web application was built to demonstrate the smooth molecule optimization of PanGu (see the Additional experiments part of the Supplementary Information section).

## Conclusions

For the conflict that large number of chemical structures exist while only a few of them being well-annotated, pre-training models can be the mediator linking the unlabeled and labeled ones, by compressing, representing and reconstructing the molecular structures, making it capable of convenient application to drug discovery downstream tasks. As far as we know, the pre-training dataset of PanGu is the largest and covers multiple public data sources. The modelling idea – MSFT was implemented by cVAE architecture to transform the graphs of small molecules into corresponding formula strings, which avoiding the difficulty of graph generation existed in graph2graph models and can provide more information during training than seq2seq models. Besides, the design of layered latent space further improved the expressive power of PanGu for both fine-tuning and fingerprinting. The novel network architecture of PanGu is easy to train and achieved the possibility that updating only one backbone but benefits for all steps of drug discovery tasks, as we evaluated the experiments above. As AI drug discovery is gradually forming the new trend, key procedures related to machine learning and data mining can be efficiently improved from accumulated data, where PanGu will be surely enhance the ability of these data. Besides molecular property prediction part of PanGu, its new chemical space discovery and automatic generation can give drug R&D scientists new inspiration and more possibility to effective experiments. We believed that PanGu will be the foothold of other new pre-training models, can efficiently promote the development of AI drug R&D and finally aid in speeding up drug discovery and raising the success rate.

## Methods

### Graph-to-sequence architecture of PanGu

PanGu is a cVAE-based graph2seq model (Fig. 4). Unlike conventional autoencoders with the same data types for both the input and output, for PanGu, we fed 2D undirected cyclic graphs as input of the encoder, obtained the encoder-derived latent spaces, and then decoded them into text-based representations as the output of the decoder. Rather than SMILES, we took advantage of a deep learning-friendly formula SELFIES as the decoder output, due to the nearly 100% validity of output chemicals without any adaptation.

**Fig. 4.**
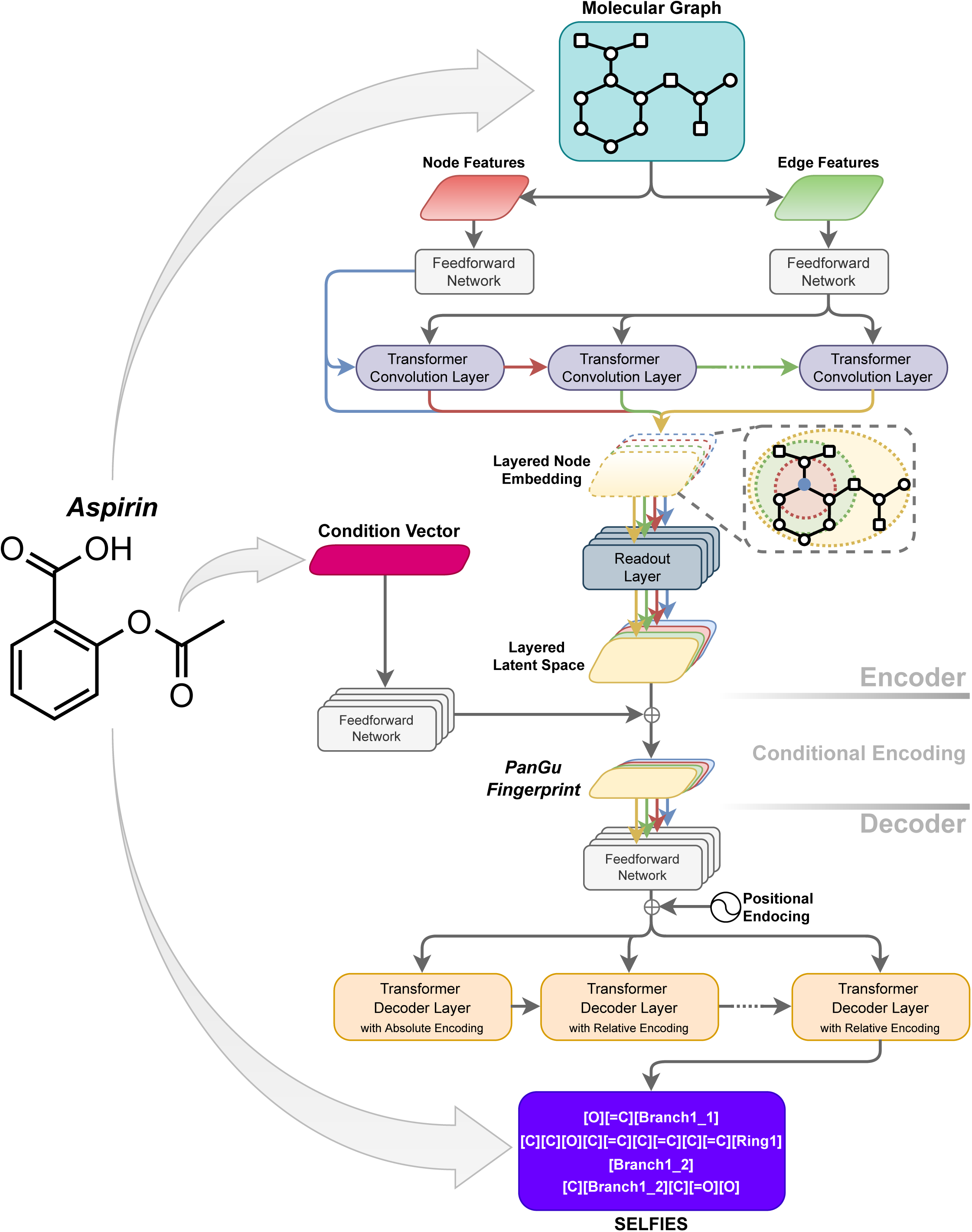
The conditional variational autoencoder architecture of PanGu. For example, the encoder of model takes molecular graph of *Aspirin* as input and output the layered latent space. Then, the layered latent space and the condition vector of *Aspirin* are integrated and input to the decoder to get the SELFIES string of *Aspirin*.

The latent spaces of the encoder can be used for unsupervised representation and supervised learning tasks. Besides, The decoder of model can generate chemicals under guidance of given molecular properties of interest in a cVAE fashion, such as molecular weight, logP, SA score and quantitative estimation of drug-likeness (QED). The molecular properties are directly incorporated into the decoder. The objective function is given by:

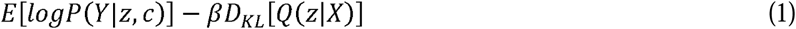

where *E* denotes an expectation value, *P* and *Q* are probability distributions, *D*_*KL*_ is the Kullback-Leibler divergence, *β* is the regularization coefficient that balances latent channel capacity and independence constraints with reconstruction accuracy, and *Y* denote molecular graphs and molecular formulas, and *z* and *c* indicate latent spaces and conditional vectors while the conditional vector is the target molecular properties and the latent space involve the molecular structure and other properties. The first and second terms are often called the reconstruction error and the KL term, respectively. In PanGu, *Q (z*|*X)* and *P (Y*|*z,c)* are approximated by an encoder and a decoder, respectively.

In the encoder part of PanGu, transformer convolution (TransformerConv)^36^ layers are used to aggregate node representation with its neighbors, and the mean-and-max readout function is used to aggregate node representations into a fixed-size graph representation. Different from the conventional VAE, we designed a layered latent space for PanGu which implemented the main idea of ECFP – describing a molecule in multiple radius scales. We concatenated multiple intermediate graph representations which passed different number of TransformerConv layers (to have different receptive fields) into a matrix as the final graph representation of a molecule, which is used as the latent space of VAE, so called layered latent space. For the decoder part, a transformer-style decoder^37^ is adopted to decompress the layered latent space into SELFIES.

### Large-scale pre-training of PanGu

We collected about 1.7 billion SMILES of molecules from ZINC20^7^, DrugSpaceX^31^ and UniChem^32^ as a large-scale pre-training data set. We split the data into three sets - training and validation sets for pre-training and model selection, and testing set for subsequent experiments. All chemicals were prepared into graphs and canonical SELFIES before training (see the Details of PanGu part of the Supplementary Information section).

PanGu was implemented and pre-trained on *MindSpore* (An open-source deep learning framework powered by Huawei) with ∼104M parameters (see the Details of PanGu part of the Supplementary Information section). While the auto-parallel module of MindSpore can automatically select a least cost model splitting strategy to achieve automatic distributed parallel training, PanGu was pre-trained on a cluster of 256 Ascend 910 AI processors for 10 epochs within 15 days.

### Detailed information of experiments

#### Molecule property prediction

We adopted scaffold splitting to split all dataset into training, validation and testing sets following an 8:1:1 ratio, same to Rong et al.’s study^38^. And we ran three independent trials with difference random seeds to get the averaged test performance. While deep learning models generally require careful hyperparameter tuning, we tried 400 random combinations of hyper-parameters and reported the best result on each task for PanGu fine-tuning. While the MLP predictor using PanGu Fingerprint or ECFP as input requires fewer hyper-parameters, therefore only 50 trials with random combinations of hyper-parameters were performed on each task.

#### Compound-protein interaction prediction

Molecules were represented by PanGu Fingerprint, and proteins were represented by a word2vec model simply with a mean-and-max pooling (the word2vec model is trained by Gensim 3.8.3 using 3-mers with 100 hidden dimensions on each dataset). Representation of molecule and protein were jointly input an MLP classifier to predict whether the CPI is active or not. In accordance with previous studies, human and *C. elegans* datasets were randomly split for 10 times with an 8:1:1 ratio for training/validation/test, and means and standard deviations of ROC-AUC, precision and recall were reported; and for the pre-processed BindingDB dataset, we reported the PRC-AUC and ROC-AUC. For each dataset, we tried 400 random combinations of hyper-parameters and reported the best result.

#### Large-scale generative database

For generating molecules, the prior distribution of latent space is obtained by using gaussian mixture model with 5 components from randomly sampled 100,000 molecules of test set. Three statistical measurements were used to evaluate the performance, they are: 1) *Validity*, which represents the proportion of generated chemically valid molecules. 2) *Uniqueness*, which represents the proportion of non-repetitive samples among the generated molecules. 3) *Novelty*, which represents the proportion of new samples among the generated molecules that are absent from the training data, indicating the ability of PanGu to generate novel molecules.

#### Constrained molecule optimization

To optimize a molecule *m*, we started from its latent representation *z* and condition vector *c*. And we randomly increase logP and decrease SA score in the condition vector *c* for 80 times to obtain 80 optimized molecules for each molecule. To systematically evaluate PanGu’s optimization capacity, we randomly sampled 50,000 molecules from the test set, and then we selected 800 starting molecules with the lowest property score from it as Jin et al.^39^ did. We reported the success rate, average property improvement of success cases and average molecular similarity between *m* and *m’*. Tanimoto similarity with Morgan fingerprint^10^ was used as the metric (radius is set to 2, according to Jin et al.’s public code).

## Data availability

The full pre-training datasets are available on ZINC20 at http://zinc20.docking.org/tranches/home/, on DrugSpaceX at https://drugspacex.simm.ac.cn/download/ and on UniChem at https://www.ebi.ac.uk/unichem/info/downloads. The datasets of molecule property prediction tasks are provided on MoleculeNet at http://moleculenet.ai/datasets-1. Three datasets of compound-protein interaction prediction are available at https://github.com/lifanchen-simm/transformerCPI.

## Author contributions

N.Q. designed and conceived the project. X.Y.L. and C.X. wrote the manuscript. X.Y.L., X.F.Z. and B.L.N performed the experiments and data analysis with help of R.Q.P. and under the guidance of N.Q., J.L.C., F.Y., C.X. and Z.P.X., N.Q. and M.Y.Z. revised the manuscript. All authors read and approved the final manuscript.

## Supplementary Information

### Details of PanGu

The encoder of PanGu was composed by 10 transformer convolution layers with 512-dimensional hidden units and 6 attention heads. Graph representations passing the 1^st^, 2^nd^, 3^rd^, 4^th^, 5^th^, 6^th^, 8^th^, 10^th^ transformer layers were read out and concatenated to obtain a latent matrix of 8×256. The decoder was consisted of 6 decoding layers – 1 with absolute positional encoding layer was followed by 5 with relative positional encoding layers, and each layer had 512-dimensional hidden units and 8 attention heads. The β of objective function was set to 0.001. The *Adam* optimizer was adopted for parameter updating and the polynomial scheduler was used for learning rate adjusting during training.

RDKit was utilized to extract features of atoms and bonds. Details of feature extraction is showed in Supplementary Table 1 and Supplementary Table 2.

### Detailed Information about molecule property prediction tasks

The details of each dataset are listed below:

#### Molecular Classification Datasets

- BBBP^40^ records whether a compound has the permeability property to penetrate the blood-brain barrier.
- SIDER^41^ records adverse drug reactions of marketed drugs.
- ClinTox^42^ records the difference comparison between FDA-approved drugs and eliminated toxic drugs during clinical trials.
- BACE^43^ records whether compounds have been reported to be inhibitors of human β-secretase 1 (BACE-1).
- Tox21^44^ is the dataset for 2014 Tox21 Data Challenge, containing the toxicity of compounds.
- ToxCast^45^ is a high-throughput screening test based dataset, containing multiple toxicity labels of compounds.

#### Molecular Regression Datasets

- ESOL^46^ is a small dataset of the solubility of compounds.
- Lipophilicity^8^ is based on ChEMBL, containing octanol/water distribution coefficient experiment based molecular membrane permeability and solubility of compounds.
- FreeSolv^47^ is a subset of Free Solvation Database, containing both experiment and alchemical free energy calculation-based hydration free energy of small molecules in water.
- QM7^48^ is a subset of GDB-13, containing the computed atomization energies of stable and synthetically accessible organic molecules, such as HOMO/LUMO, atomization energy and so on for various molecular structures such as triple bonds, cycles, amide, epoxy, etc.
- QM8^49^ contains computer-generated quantum mechanical properties, e.g., electronic spectra and excited state energy of small molecules.

### Ablation studies of layered latent space

We compared PanGu with or without layered latent space to verify the difference of expressive power. Two versions of model have nearly the same number of parameters (∼55M parameters) and both pre-trained by using ZINC250k dataset from scratch. Furthermore, we evaluated both of them on 8 molecule property prediction tasks with same strategy to the Molecule property prediction part of the Methods section. The training and validation losses are showed in Supplementary Figure 1, PanGu with layered latent space outperformed in both training and validation. And the result of molecule property prediction tasks shown in Supplementary Table 3 also proved the high expressive power of layered latent space.

### Additional experiments

#### Molecule validity and reconstruction

This task is applied to test PanGu on the decoding validity when latent spaces are sampled from prior, and the reconstruction accuracy of molecules from the test set. These two measurements are closely related to the pre-training task and can directly reflect the performance of the model on the pre-training task. While pre-training models i.e. GROVER and X-Mol are not VAE-based, therefore we compared PanGu to multiple VAE-based models, they are JT-VAE^39^, CharVAE^50^, GrammerVAE^33^, SD-VAE^51^ and GraphVAE^52^. To compute validity of molecule sampling, we randomly sampled 1000 latent vectors from the standard normal distribution as prior, and decoded these vectors into molecules. The percentage of chemically valid molecules among the results is calculated as validity. For the molecule reconstruction, since the encoding process is stochastic, we followed the same strategy of Kusner et al.^33^ which is scaling the standard deviation of the latent distribution *Q* (*z*|*X)* by 0.01 to produce more consistent reconstruction results. To obtain the reconstruction accuracy, we randomly sampled 5,000 molecules from the test set. Each molecule was reconstructed 100 times to get 100 molecules. The portion of the reconstructed molecules that are identical to the input was computed for each molecule. And finally, the average of portions was reported as reconstruction accuracy. The results showed that PanGu achieved 100% molecule sampling validity and 91.4% reconstruction accuracy (Supplementary Table 4), outstandingly outperforming all other six models: by 14.7% improvement in reconstruction accuracy compared to the second-ranking model JT-VAE and tied for the first place with JT-VAE in validity. These results demonstrated that the large-scale pre-training of MSFT enables PanGu to learn the molecular representation quite well.

#### Molecule property prediction with random split

Here, datasets were randomly split into training, validation and testing sets with an 8:1:1 ratio and took DMPNN as baseline. While the original data-splits of Yang et al.’s study were hard to reproduce, we evaluated PanGu on 10 different trials to get the averaged, median and best test performance. Also, we only tried 50 and 10 random combinations of hyper-parameters on each task for fine-tuning and fingerprinting respectively to get competitive results. The results were shown in Supplementary Table 5.

#### Drug-drug interaction prediction

A drug-drug interaction (DDI) can be defined as an interaction between drugs that prevents the drug from performing as expected, including some unknown mechanisms of adverse drug events (ADEs); therefore, DDI prediction has become another important task related to in silico molecular analysis. The dataset of this task includes 192,284 DDIs and 86 DDI types with preprocessed training, validation, and testing sets of 115,446, 38,419 and 38,419 DDIs, respectively. The prediction of drug-drug interaction is a multi-classification task which inputs chemical structures of two drugs and outputs the DDI types. PanGu Fingerprint was used to represent each molecule. Firstly, molecular fingerprints of a DDI pair were projected through the same feedforward network and summed up element-wisely to get the representation of the DDI pair, then it passed through another feedforward network of a predictor to obtain the probabilities of each DDI type as output. For the pre-processed dataset, we used the validation set for model selection, and reported the accuracy on test set. Total 400 random combinations of hyper-parameters were tried to get the best result. X-Mol (fine-tuned)^15^ and DeepDDI^53^ were compared in this task. The results showed that PanGu Fingerprint obtained an accuracy of 0.957, which is higher than the values of 0.952 and 0.924 achieved by X-Mol and DeepDDI, respectively (Supplementary Figure 2a).

#### Chemical reaction productivity prediction

The prediction of chemical reaction productivity is a necessary computational task in the field of compound production and drug synthesis. The prediction of this task should consider base, ligand, aryl halide and additive compounds that participate in or affect the chemical reaction. The dataset of this task contains 3,956 chemical reactions and corresponding productivity value. PanGu Fingerprint was used to represent four parts of the chemical reaction - reactants, reaction environment, catalysts and other components that participate the reaction. Four fingerprints of a reaction were concatenated and used for making prediction by a feedforward predictor network. The output was productivity value. The dataset was split into training and test data with a ratio of 7:3, and the 10-fold cross-validation was adopted as previous studies did and the average of RMSEs was reported. Total 400 random combinations of hyper-parameters were tried to get the best result. The result was compared to X-Mol (fine-tuned)^15^ and Random Forest^54^. As a result, PanGu Fingerprint obtained an average RMSE of 5.82, which is significantly lower than the values of 6.29 and 7.80 reported by X-Mol and the Random Forest, respectively (Supplementary Figure 2b).

#### Small-scale molecule generation

Six generative models were compared in this task, i.e.: JT-VAE, GCPN^55^, MRNN^56^, GraphNVP^57^, GraphAF^58^ and X-Mol. In order to get a consistent comparison, ZINC250K dataset was adopted to pre-train the PanGu from scratch. In the generation phase, gaussian mixture models (with 3 components) were used to fit each dimension of latent space in training set as prior. Then we sample 10,000 latent spaces from the prior distribution and feed into decoder to get 10,000 generated molecules for evaluation. As a result, PanGu outperformed other models with 100% on validity, uniqueness and novelty measurements (Supplementary Table 6).

#### Smooth molecule optimization using web application

We built a simple web application of molecule optimization based on Flask, which allows us to input a SMILES of molecule and modify properties smoothly by dragging sliders to get the optimized one (Supplementary Figure 3). To test the web application, we selected a base molecule from the test set which has moderate properties, and them we performed modification on each property independently by turning up or tuning down. Finally, we record 4 optimized molecules for each property, two of them have lower property than the original one and the other two are higher. The result clearly demonstrated the ability of PanGu for smooth molecule optimization (Supplementary Figure 4). We found that the difference of property between optimized molecule and original molecule is inversely proportional to the similarity. It’s easy to understand that the property difference usually means the degree of modification, and the more modifications the similarity decreases. But of course, there is an exception. In case of QED (Supplementary Figure 3d), the molecule with the highest property (QED=0.8665) is much more similar to the original molecule (QED=0.7438) than the molecule with second-high property (QED=0.8265), the similarities are 0.4118 and 0.3738, respectively.

## Data availability

The datasets of drug-drug interaction prediction and chemical reaction productivity prediction are available at https://bitbucket.org/kaistsystemsbiology/deepddi/src/master/data/ and https://github.com/doylelab/rxnpredict, respectively.

## Supplementary Figure Legends

**Supplementary Figure 1.**
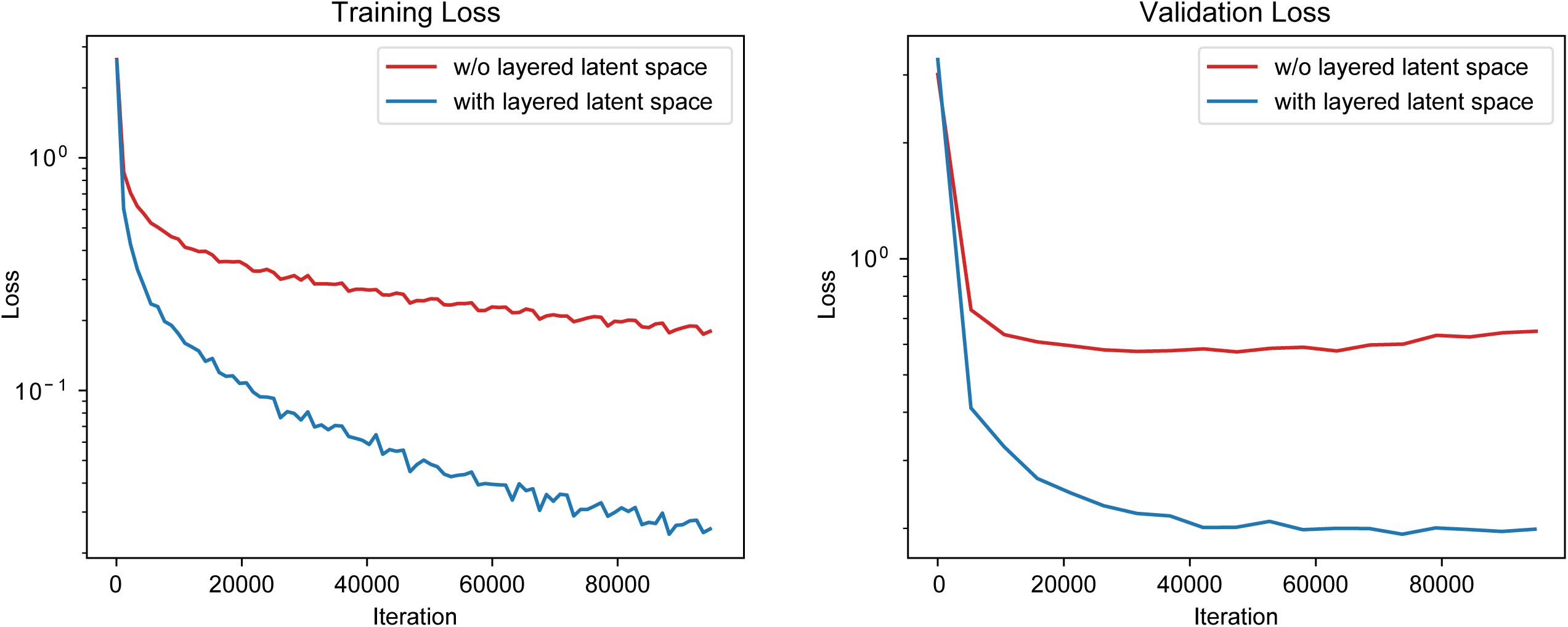
Training and validation loss of PanGu with or without layered latent space.

**Supplementary Figure 2.**
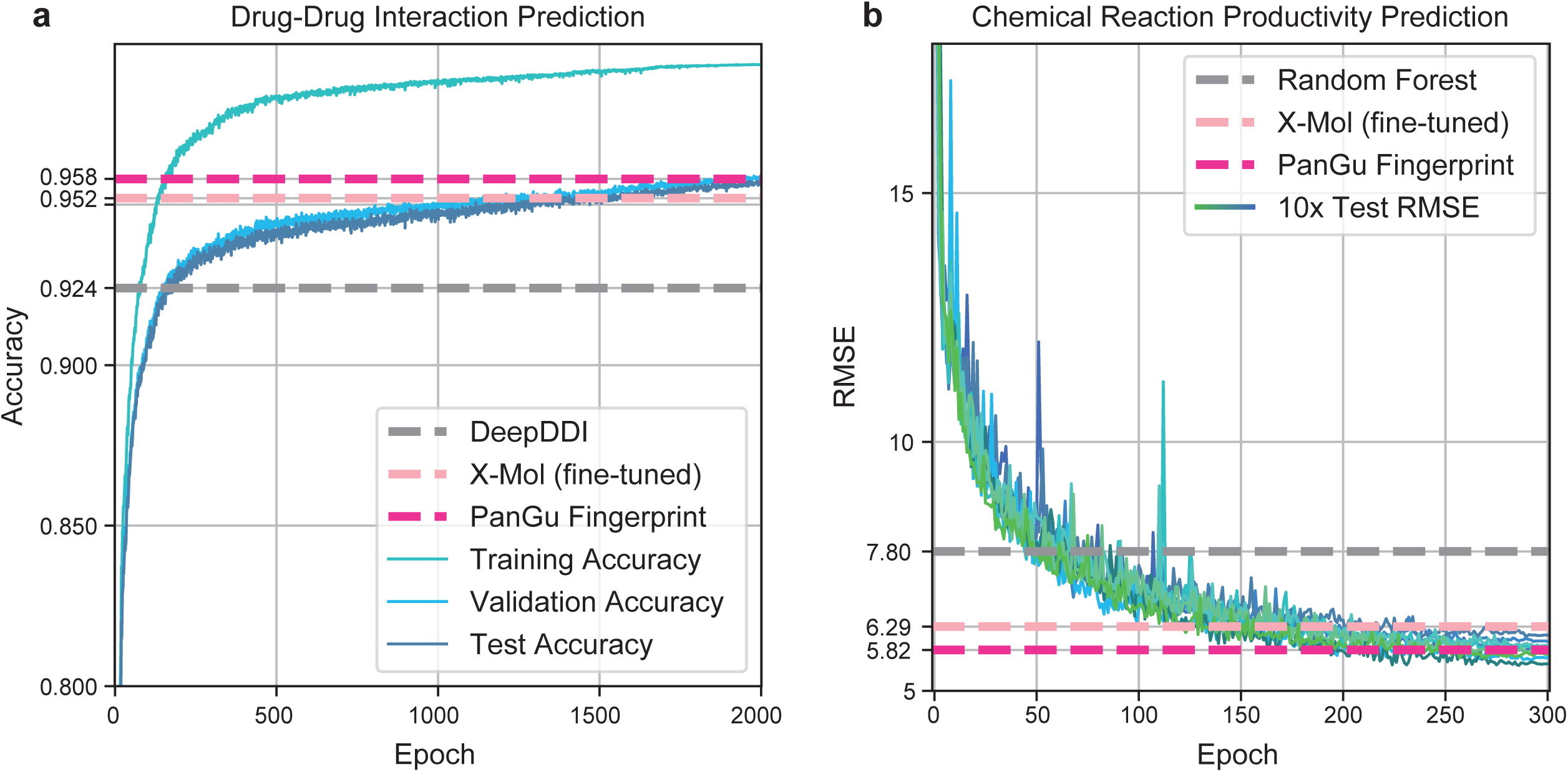
Performance comparison of PanGu Fingerprint with baseline on drug-drug prediction and chemical reaction productivity prediction. **a**, The epoch-accuracy curves of PanGu Fingerprint on training, validation and test data of the drug-drug interaction prediction task. Horizontal dashed lines indicate the test accuracy of DeepDDI, X-Mol (fine-tuned) and PanGu Fingerprint. **b**, The epoch-test root mean square error (RMSE) curves of PanGu Fingerprint under 10-fold cross validation for the chemical reaction productivity prediction. Horizontal dashed lines indicate the test RMSE of random forest, X-Mol (fine-tuned) and PanGu Fingerprint.

**Supplementary Figure 3.**
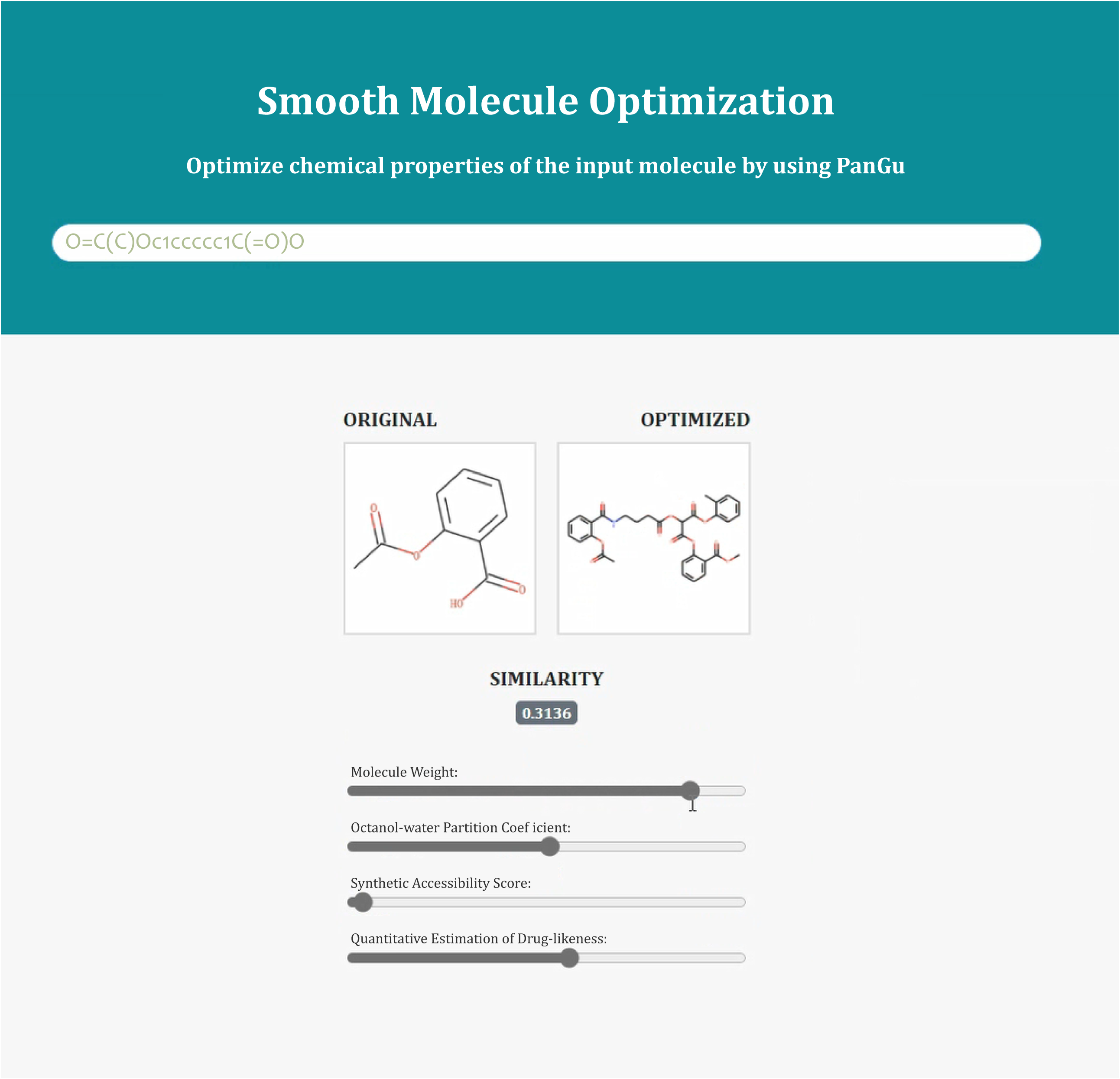
Molecule optimization web application. In the case of figure, we input Aspirin and drag the slider of molecule weight to obtain a larger molecule, and we can see that the optimized molecule contains the entire structure of the original one.

**Supplementary Figure 4.**
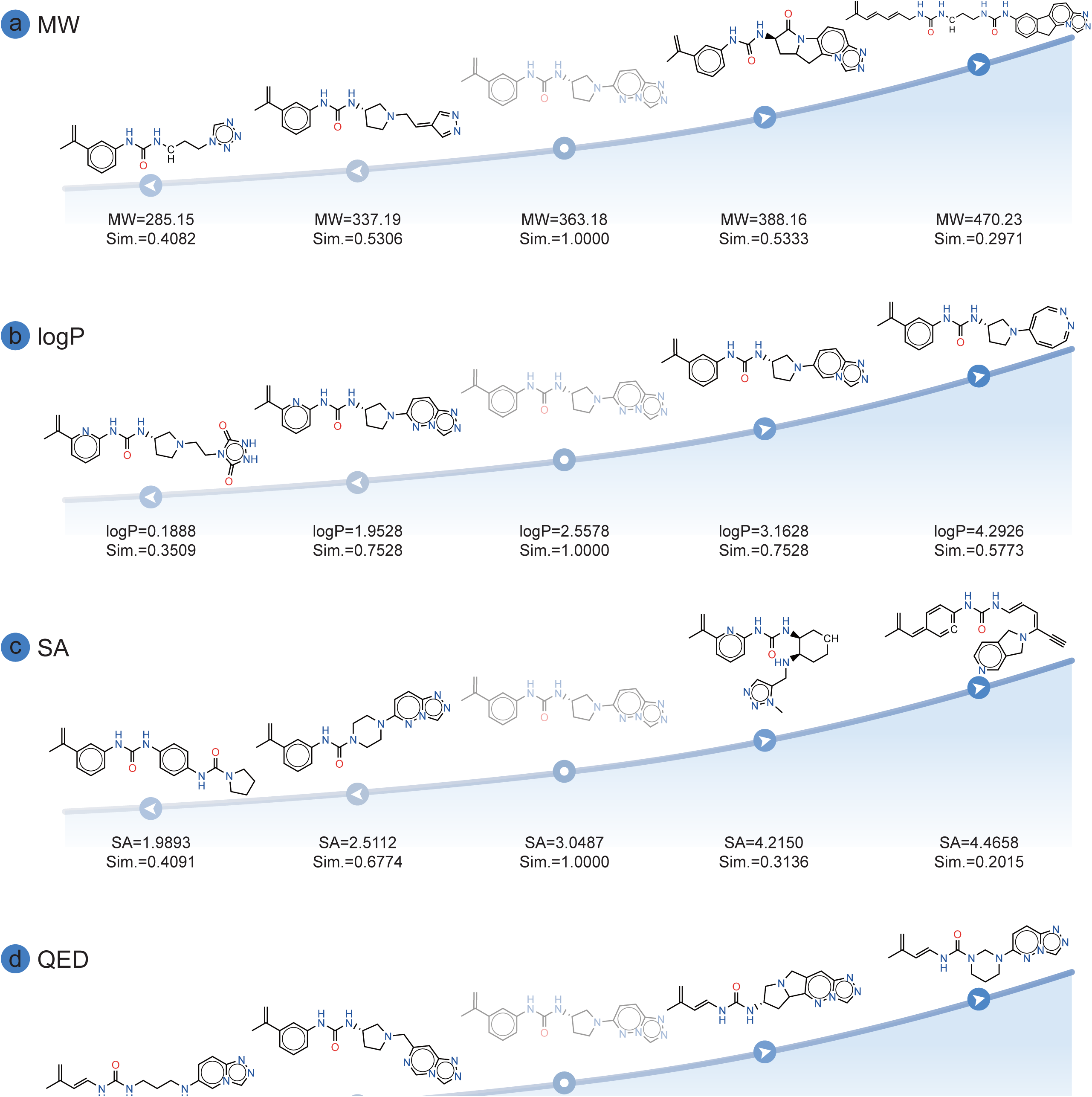
Smooth optimization using PanGu by turning-up and -down each property independently. The starting molecule with moderate properties was place at the middle row of figure.

**Supplementary Figure 5.**
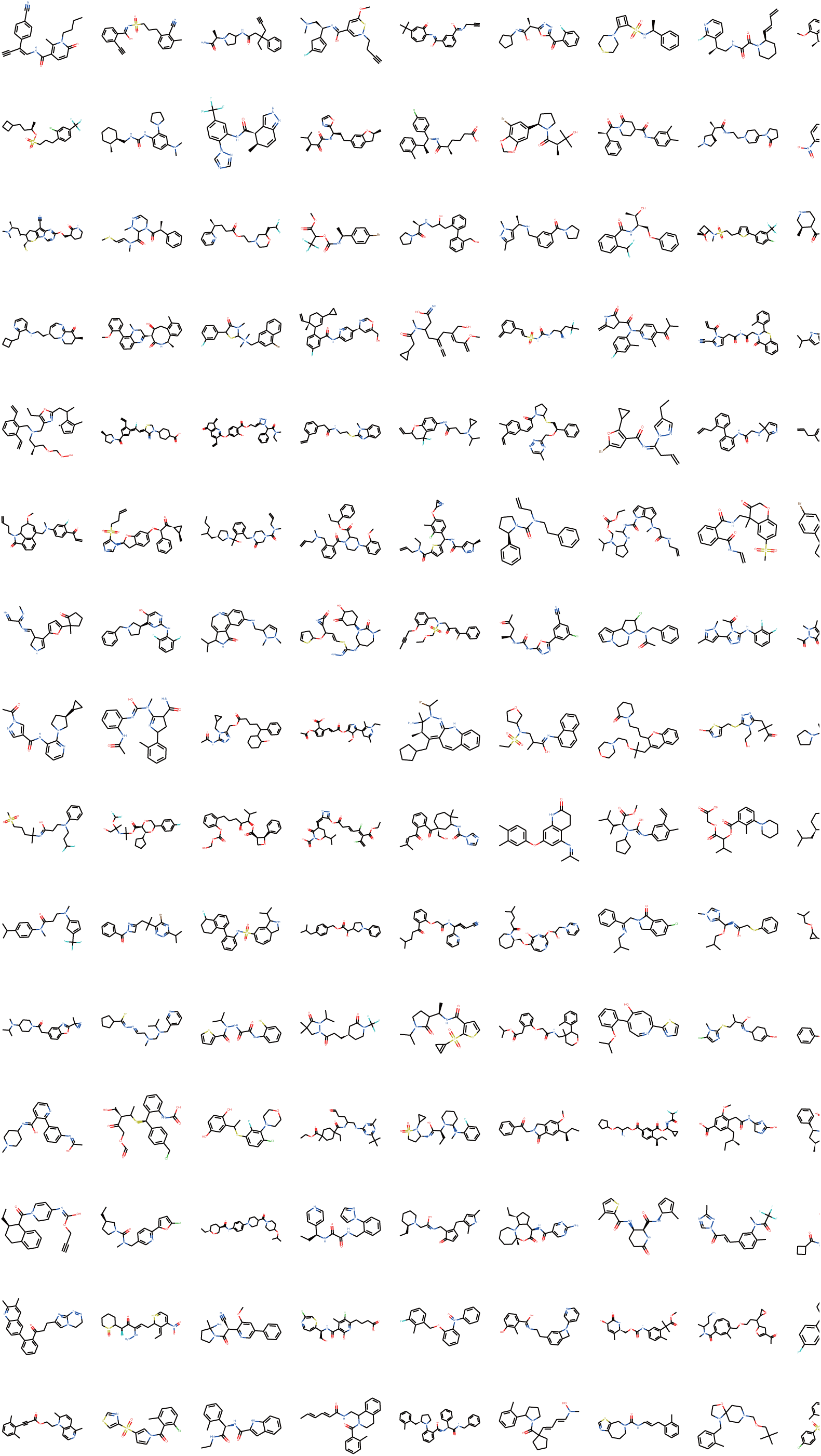
150 molecules sampled from 100 million PanGu generated set.

## Supplementary Tables

**Supplementary Table 1.** Atom features.

| Feature | Size | Description |
| --- | --- | --- |
| Atom Type | 43 | [B, C, N, O, F, Si, P, S, Cl, As, Se, Br, Te, I, At, ...] (one-hot) |
| Atomic Mass | 1 | mass of the atom, divided by 100 (float) |
| Aromaticity | 1 | whether the atom is part of an aromatic system [0/1] (one-hot) |
| Number of Bonds | 9 | number of covalent bonds [0, 1, 2, 3, 4, 5, 6, 7, 8] (one-hot) |
| Number of H | 5 | number of connected hydrogens [0, 1, 2, 3, 4] (one-hot) |
| Formal Charge | 1 | electrical charge (integer) |
| Radical Electrons | 1 | number of radical electrons (integer) |
| Chirality | 3 | [none, R, S] (one-hot) |
| Hybridization | 6 | [sp, sp2, sp3, sp3d, sp3d2, other] (one-hot) |

**Supplementary Table 2.** Bond features.

| Feature | Size | Description |
| --- | --- | --- |
| Bond Type | 4 | [single, double, triple, aromatic] (one-hot) |
| Conjugation | 1 | whether the bond is conjugated [0/1] (one-hot) |
| Ring | 1 | whether the bond is in ring [0/1] (one-hot) |
| Stereo | 6 | [None, Any, Z, E, Cis, Trans] (one-hot) |

**Supplementary Table 3.** Ablation study: comparison between PanGu Fingerprint with and without layered latent space on multiple molecule property prediction tasks. Three trials with randomly seeded scaffold splitting were performed on each dataset, and the mean and standard deviation of the performance values were reported.

| Task | Metric | PanGu Fingerprint<br>w/o layered latent space | PanGu Fingerprint<br>with layered latent space |
| --- | --- | --- | --- |
| BBBP | ROC-AUC | 0.907 (0.030) | **0.915 (0.022)** |
| BACE | ROC-AUC | 0.887 (0.022) | **0.890 (0.017)** |
| Tox21 | ROC-AUC | 0.782 (0.036) | **0.816 (0.017)** |
| ToxCast | ROC-AUC | 0.685 (0.011) | **0.700 (0.004)** |
| FreeSolv | RMSE | 2.610 (0.255) | **1.755 (0.230)** |
| ESOL | RMSE | 0.869 (0.019) | **0.818 (0.034)** |
| Lipophilicity | RMSE | 0.814 (0.007) | **0.708 (0.062)** |
| QM7 | MAE | 114.7 (7.2) | **91.4 (3.2)** |

The bold indicates the cases of the models outperforming another.

**Supplementary Table 4.** Prior validity and reconstruction accuracy results.

| Method | Architecture | Prior Validity | Reconstruction Accuracy |
| --- | --- | --- | --- |
| CharVAE | VAE | 0.7% | 44.6% |
| GrammarVAE | VAE | 7.2% | 53.7% |
| SD-VAE | VAE | 43.5% | 76.2% |
| GraphVAE | VAE | 13.5% | - |
| JT-VAE | VAE | **100.0%** | 76.7% |
| PanGu<sup>*</sup> | VAE | **100.0%** | **91.4%** |

The results marked with <sup>\*</sup> are provided by this study, and baseline results are copied from Jin et al. (2018).
And the bold indicates the cases of the models outperforming other models.

**Supplementary Table 5.** Comparison between PanGu and DMPNN on multiple molecule property prediction tasks. Totally 10 trials with random splitting were performed on each dataset, and the mean, standard deviation, median and best of the performance values were reported.

| Task | Metric | DMPNN | Fine-tuned PanGu |  |  | PanGu Fingerprint |  |  |
| --- | --- | --- | --- | --- | --- | --- | --- | --- |
|  |  |  | 10x Mean | Median | Best | 10x Mean | Median | Best |
| BBBP | ROC-AUC | 0.913 | **0.9244**<sub>(0.0181)</sub> | 0.9282 | 0.9481 | 0.9186<sub>(0.0199)</sub> | 0.9114 | 0.9506 |
| SIDER | ROC-AUC | 0.646 | **0.6483**<sub>(0.0248)</sub> | 0.6580 | 0.6790 | 0.6414<sub>(0.0270)</sub> | 0.6430 | 0.6760 |
| ClinTox | ROC-AUC | 0.894 | **0.8958**<sub>(0.0538)</sub> | 0.9125 | 0.9393 | 0.8884<sub>(0.0325)</sub> | 0.8900 | 0.9262 |
| BACE | ROC-AUC | 0.878 | 0.8895<sub>(0.0192)</sub> | 0.8917 | 0.9146 | **0.8898**<sub>(0.0336)</sub> | 0.8776 | 0.9431 |
| Tox21 | ROC-AUC | 0.845 | **0.8506**<sub>(0.0132)</sub> | 0.8486 | 0.8719 | 0.8375<sub>(0.0158)</sub> | 0.8360 | 0.8679 |
| ToxCast | ROC-AUC | 0.737 | **0.7652**<sub>(0.0105)</sub> | 0.7645 | 0.7795 | 0.7455<sub>(0.0084)</sub> | 0.7446 | 0.7632 |
| HIV | ROC-AUC | 0.816 | **0.8324**<sub>(0.0170)</sub> | 0.8260 | 0.8640 | 0.8158<sub>(0.0346)</sub> | 0.8168 | 0.8791 |
| FreeSolv | RMSE | 1.167 | **0.9522**<sub>(0.2473)</sub> | 0.9266 | 0.4962 | 1.0689<sub>(0.1481)</sub> | 1.0845 | 0.8098 |
| ESOL | RMSE | 0.665 | **0.5747**<sub>(0.0365)</sub> | 0.5847 | 0.4785 | 0.6039<sub>(0.595)</sub> | 0.6020 | 0.5093 |
| Lipophilicity | RMSE | 0.596 | **0.5586**<sub>(0.0331)</sub> | 0.5538 | 0.5041 | 0.6811<sub>(0.0260)</sub> | 0.6786 | 0.6445 |
| QM7 | MAE | 66.475 | **60.922**<sub>(4.0387)</sub> | 60.844 | 53.962 | 63.800<sub>(4.0715)</sub> | 63.613 | 57.421 |
| QM8 | MAE | 0.011 | **0.0085**<sub>(0.0002)</sub> | 0.0086 | 0.0080 | 0.0146<sub>(0.0002)</sub> | 0.0147 | 0.0144 |

The results of DMPNN are copied from Yang et al. (2019).

**Supplementary Table 6.** Small-scale molecule generation results. *Validity w/o check* is only evaluated on models with valency constraints.

| Method | Type of Generation | Validity | Validity w/o check | Uniqueness | Novelty |
| --- | --- | --- | --- | --- | --- |
| GraphNVP<sup>a</sup> | Graph-based | 100.00% | - | 94.80% | 100.0% |
| JT-VAE<sup>a</sup> | Graph-based | 100.00% | - | 100.0% | 100.0% |
| GCPN<sup>a</sup> | Graph-based | 100.00% | 20.00% | 99.97% | 100.0% |
| MRNN<sup>a</sup> | Graph-based | 100.00% | 65.00% | 99.89% | 100.0% |
| GraphAF<sup>a</sup> | Graph-based | 100.00% | 68.00% | 99.10% | 100.0% |
| X-Mol<sup>b</sup> | Sequence-based | 85.28% | - | 99.91% | 100.0% |
| PanGu<sup>*</sup> | Sequence-based | 100.00% | - | 100.0% | 100.0% |

<sup>\*</sup> The results are provided by this study. <sup>a</sup> The results are taken from Shi et al. (2020). <sup>b</sup> The results are taken from Xue et al. (2020).

## Reference

1. World Health Organization. International Classification of Diseases (ICD). http://www.who.int/classifications/icd/en/.

2. ProductDevelopment, I. of M. (US) C. on A. R. D. R. and O., Field, M. J. & Boat, T. F. Profile of Rare Diseases. Rare Diseases and Orphan Products: Accelerating Research and Development (National Academies Press (US), 2010).

3. Sertkaya, A., Wong, H.-H., Jessup, A. & Beleche, T. Key cost drivers of pharmaceutical clinical trials in the United States. Clin. Trials Lond. Engl. 13, 117–126 (2016).

4. Burki, T. A new paradigm for drug development. Lancet Digit. Health 2, e226–e227 (2020).

5. Zhavoronkov, A. et al. Deep learning enables rapid identification of potent DDR1 kinase inhibitors. Nat. Biotechnol. 37, 1038–1040 (2019).

6. Kapetanovic, I. M. Computer-aided drug discovery and development (CADDD): In silico-chemico-biological approach. Front. Pharmacol. Toxicol. 171, 165–176 (2008).

7. Irwin, J. J. et al. ZINC20—A Free Ultralarge-Scale Chemical Database for Ligand Discovery. J. Chem. Inf. Model. 60, 6065–6073 (2020).

8. Gaulton, A. et al. ChEMBL: a large-scale bioactivity database for drug discovery. Nucleic Acids Res. 40, D1100–D1107 (2012).

9. Kim, S. et al. PubChem Substance and Compound databases. Nucleic Acids Res. 44, D1202–D1213 (2016).

10. Rogers, D. & Hahn, M. Extended-Connectivity Fingerprints. J. Chem. Inf. Model. 50, 742–754 (2010).

11. Lim, J., Ryu, S., Kim, J. W. & Kim, W. Y. Molecular generative model based on conditional variational autoencoder for de novo molecular design. J. Cheminformatics 10, 31 (2018).

12. Kearnes, S., McCloskey, K., Berndl, M., Pande, V. & Riley, P. Molecular graph convolutions: moving beyond fingerprints. J. Comput. Aided Mol. Des. 30, 595–608 (2016).

13. Kwon, Y. et al. Efficient learning of non-autoregressive graph variational autoencoders for molecular graph generation. J. Cheminformatics 11, 70 (2019).

14. Rong, Y. et al. Self-Supervised Graph Transformer on Large-Scale Molecular Data. Adv. Neural Inf. Process. Syst. 33, 12559–12571 (2020).

15. Xue, D. et al. X-MOL: large-scale pre-training for molecular understanding and diverse molecular analysis. bioRxiv 2020.12.23.424259 (2020) doi:10.1101/2020.12.23.424259.

16. Weininger, D. SMILES, a chemical language and information system. 1. Introduction to methodology and encoding rules. J. Chem. Inf. Comput. Sci. 28, 31–36 (1988).

17. Hodgson, J. ADMET—turning chemicals into drugs. Nat. Biotechnol. 19, 722–726 (2001).

18. Wu, Z. et al. MoleculeNet: a benchmark for molecular machine learning. Chem. Sci. 9, 513–530 (2018).

19. Kipf, T. N. & Welling, M. Semi-Supervised Classification with Graph Convolutional Networks. 160902907 Cs Stat (2017).

20. Schütt, K. et al. SchNet: A continuous-filter convolutional neural network for modeling quantum interactions. in Advances in Neural Information Processing Systems vol. 30 (2017).

21. Gilmer, J., Schoenholz, S. S., Riley, P. F., Vinyals, O. & Dahl, G. E. Neural message passing for Quantum chemistry. in Proceedings of the 34th International Conference on Machine Learning - Volume 70 1263–1272 (http://JMLR.org, 2017).

22. Yang, K. et al. Analyzing Learned Molecular Representations for Property Prediction. J. Chem. Inf. Model. 59, 3370–3388 (2019).

23. Xiong, Z. et al. Pushing the Boundaries of Molecular Representation for Drug Discovery with the Graph Attention Mechanism. J. Med. Chem. 63, 8749–8760 (2020).

24. RDKit: Open-source Cheminformatics (RDKit); http://www.rdkit.org.

25. Vamathevan, J. et al. Applications of machine learning in drug discovery and development. Nat. Rev. Drug Discov. 18, 463–477 (2019).

26. Liu, H., Sun, J., Guan, J., Zheng, J. & Zhou, S. Improving compound-protein interaction prediction by building up highly credible negative samples. Bioinforma. Oxf. Engl. 31, i221–229 (2015).

27. Tsubaki, M., Tomii, K. & Sese, J. Compound-protein interaction prediction with end-to-end learning of neural networks for graphs and sequences. Bioinforma. Oxf. Engl. 35, 309–318 (2019).

28. Gao, K. Y. et al. Interpretable Drug Target Prediction Using Deep Neural Representation. 3371–3377 (2018).

29. Chen, L. et al. TransformerCPI: improving compound-protein interaction prediction by sequence-based deep learning with self-attention mechanism and label reversal experiments. Bioinforma. Oxf. Engl. 36, 4406–4414 (2020).

30. Nguyen, T. et al. GraphDTA: Predicting drug–target binding affinity with graph neural networks. bioRxiv 684662 (2020) doi:10.1101/684662.

31. Yang, T. et al. DrugSpaceX: a large screenable and synthetically tractable database extending drug space. Nucleic Acids Res. 49, D1170–D1178 (2021).

32. Chambers, J. et al. UniChem: a unified chemical structure cross-referencing and identifier tracking system. J. Cheminformatics 5, 3 (2013).

33. Kusner, M. J., Paige, B. & Hernández-Lobato, J. M. Grammar Variational Autoencoder. in International Conference on Machine Learning 1945–1954 (PMLR, 2017).

34. Krenn, M., Häse, F., Nigam, A., Friederich, P. & Aspuru-Guzik, A. Self-referencing embedded strings (SELFIES): A 100% robust molecular string representation. Mach. Learn. Sci. Technol. 1, 045024 (2020).

35. Besnard, J. et al. Automated design of ligands to polypharmacological profiles. Nature 492, 215–220 (2012).

36. Shi, Y. et al. Masked Label Prediction: Unified Message Passing Model for Semi-Supervised Classification. 200903509 Cs Stat (2021).

37. Vaswani, A. et al. Attention Is All You Need. 170603762 Cs (2017).

38. Rong, Y. et al. Self-Supervised Graph Transformer on Large-Scale Molecular Data. 200702835 Cs Q-Bio (2020).

39. Jin, W., Barzilay, R. & Jaakkola, T. Junction Tree Variational Autoencoder for Molecular Graph Generation. in International Conference on Machine Learning 2323–2332 (PMLR, 2018).

40. Martins, I. F., Teixeira, A. L., Pinheiro, L. & Falcao, A. O. A Bayesian Approach to in Silico Blood-Brain Barrier Penetration Modeling. J. Chem. Inf. Model. 52, 1686–1697 (2012).

41. Kuhn, M., Letunic, I., Jensen, L. J. & Bork, P. The SIDER database of drugs and side effects. Nucleic Acids Res. 44, D1075–D1079 (2016).

42. Gayvert, K. M., Madhukar, N. S. & Elemento, O. A Data-Driven Approach to Predicting Successes and Failures of Clinical Trials. Cell Chem. Biol. 23, 1294–1301 (2016).

43. Subramanian, G., Ramsundar, B., Pande, V. & Denny, R. A. Computational Modeling of β-Secretase 1 (BACE-1) Inhibitors Using Ligand Based Approaches. J. Chem. Inf. Model. 56, 1936–1949 (2016).

44. Tox21 challenge, 2017. https://tripod.nih.gov/tox21/challenge/.

45. Richard, A. M. et al. ToxCast Chemical Landscape: Paving the Road to 21st Century Toxicology. Chem. Res. Toxicol. 29, 1225–1251 (2016).

46. Delaney, J. S. ESOL:L Estimating Aqueous Solubility Directly from Molecular Structure. J. Chem. Inf. Comput. Sci. 44, 1000–1005 (2004).

47. Mobley, D. L. & Guthrie, J. P. FreeSolv: a database of experimental and calculated hydration free energies, with input files. J. Comput. Aided Mol. Des. 28, 711–720 (2014).

48. Blum, L. C. & Reymond, J.-L. 970 Million Druglike Small Molecules for Virtual Screening in the Chemical Universe Database GDB-13. J. Am. Chem. Soc. 131, 8732–8733 (2009).

49. Ramakrishnan, R., Hartmann, M., Tapavicza, E. & von Lilienfeld, O. A. Electronic spectra from TDDFT and machine learning in chemical space. J. Chem. Phys. 143, 084111 (2015).

50. R, G.-B. et al. Automatic Chemical Design Using a Data-Driven Continuous Representation of Molecules. X 4, 268–276 (2018).

51. Dai, H., Tian, Y., Dai, B., Skiena, S. & Song, L. Syntax-Directed Variational Autoencoder for Structured Data. 180208786 Cs (2018).

52. Simonovsky, M. & Komodakis, N. GraphVAE: Towards Generation of Small Graphs Using Variational Autoencoders. in Artificial Neural Networks and Machine Learning – ICANN 2018 (eds. Kůrková, V., Manolopoulos, Y., Hammer, B., Iliadis, L. & Maglogiannis, I.) 412–422 (Springer International Publishing, 2018). doi:10.1007/978-3-030-01418-6_41.

53. Ryu, J. Y., Kim, H. U. & Lee, S. Y. Deep learning improves prediction of drug–drug and drug–food interactions. Proc. Natl. Acad. Sci. 115, E4304 (2018).

54. Ahneman, D. T., Estrada, J. G., Lin, S., Dreher, S. D. & Doyle, A. G. Predicting reaction performance in C–N cross-coupling using machine learning. Science 360, 186 (2018).

55. You, J., Liu, B., Ying, Z., Pande, V. & Leskovec, J. Graph Convolutional Policy Network for Goal-Directed Molecular Graph Generation. in Advances in Neural Information Processing Systems (eds. Bengio, S. et al.) vol. 31 (Curran Associates, Inc., 2018).

56. Popova, M., Shvets, M., Oliva, J. & Isayev, O. MolecularRNN: Generating realistic molecular graphs with optimized properties. 190513372 Cs Q-Bio Stat (2019).

57. Madhawa, K., Ishiguro, K., Nakago, K. & Abe, M. GraphNVP: An Invertible Flow Model for Generating Molecular Graphs. 190511600 Cs Stat (2019).

58. Shi, C. et al. GraphAF: a Flow-based Autoregressive Model for Molecular Graph Generation. 200109382 Cs Stat (2020).

